# Large scale genomic analysis shows no evidence for repeated pathogen adaptation during the invasive phase of bacterial meningitis in humans

**DOI:** 10.1101/070045

**Authors:** John A. Lees, Philip H.C. Kremer, Ana S. Manso, Nicholas J. Croucher, Bart Ferwerda, Mercedes Valls Serón, Marco R. Oggioni, Julian Parkhill, Matthjis C. Brouwer, Arie van der Ende, Diederik van de Beek, Stephen D. Bentley

## Abstract

Recent studies have provided evidence for rapid pathogen genome variation, some of which could potentially affect the course of disease. We have previously detected such variation by comparing isolates infecting the blood and cerebrospinal fluid (CSF) of a single patient during a case of bacterial meningitis.

To determine whether the observed variation repeatedly occurs in cases of disease, we performed whole genome sequencing of paired isolates from blood and CSF of 938 meningitis patients. We also applied the same techniques to 54 paired isolates from the nasopharynx and CSF.

Using a combination of reference-free variant calling approaches we show that no genetic adaptation occurs in the invasive phase of bacterial meningitis for four major pathogen species: *Streptococcus pneumoniae, Neisseria meningitidis, Listeria monocytogenes and Haemophilus influenzae*. From nasopharynx to CSF, no adaptation was seen in *S. pneumoniae*, but in *N. meningitidis* mutations potentially mediating adaptation to the invasive niche were occasionally observed in the *dca* gene.

This study therefore shows that the bacteria capable of causing meningitis are already able to do this upon entering the blood, and no further sequence change is necessary to cross the blood-brain barrier. The variation discovered from nasopharyngeal isolates suggest that larger studies comparing carriage and invasion may help determine the likely mechanisms of invasiveness.

**Author Summary:** We have analysed the entire DNA sequence from bacterial pathogen isolates from cases of meningitis in 938 Dutch adults, focusing on comparing pairs of isolates from the patient’s blood and their cerebrospinal fluid. Previous research has been on only a single patient, but showed possible signs of adaptation to treatment within the host over the course of a single case of disease.

By sequencing many more such paired samples, and including four different bacterial species, we were able to determine that adaptation of the pathogen does not occur after bloodstream invasion during bacterial meningitis.

We also analysed 54 pairs of isolates from pre- and post-invasive niches from the same patient. In *N. meningitidis* we found variation in the sequence of one gene which appears to provide bacteria with an advantage after invasion of the bloodstream.

Overall, our findings indicate that evolution after invasion in bacterial meningitis is not a major contribution to disease pathogenesis. Future studies should involve more extensive sampling between the carriage and disease niches, or on variation of the host.

## Introduction

Bacterial meningitis is a severe inflammation of the meninges surrounding the brain as a response to the presence of bacteria [1]. This inflammation can compromise brain function, requiring immediate admission to hospital. In European countries, the four bacteria which most frequently cause meningitis are *Streptococcus pneumoniae, Neisseria meningitidis, Haemophilus influenzae* and *Listeria monocytogenes* [2].

The route of infection varies depending on the species of bacteria, though in the majority of invasive cases the final stage is from blood to cerebrospinal fluid (CSF) [1]. Respiratory pathogens (*S. pneumoniae, N. meningitidis* and *H. influenzae*) are carried asymptomatically in the nasopharynx by a proportion of the population at a given time [3, 4]. *L. monocytogenes* is food-borne infection and can result from consumption of contaminated products [5, 6]. In a small number of cases commensal nasopharyngeal or ingested food-borne bacteria may invade the blood through a single cell bottleneck (bacteraemia) [7], then cross the blood-brain barrier into the CSF where they cause meningitis [8]. In some meningitis patients the CSF may be invaded directly due to CSF leakage or otitis media [9], in which case the progression of bacteria after carriage is reversed: CSF to blood.

Until recently it was thought that mutation rates in bacterial genomes were low, and as such would not change within a single host [10]. However, many studies sequencing bacterial populations of various different species gave estimates three orders of magnitude higher than previously expected [11-13]. These new estimates of mutation rate also gave evidence that DNA sequence variation can occur over the course of a single infection [14].

Such within-host variation has been shown to occur through a variety of mechanisms such as recombination [15], gene loss [16, 17] and variation in regulatory regions [18-20]. The rapid variation that occurs in these regions of the genome can increase the population’s fitness as the bacteria adapt to the host environment [21, 22], and potentially affect the course of disease [23]. Previous studies have shown variation between strains even during the rapid clinical progression of bacterial meningitis [24, 25].

It is possible that the existing genetics of the bacterium invading from the carriage population may determine, prior to blood stream invasion, whether CSF invasion is possible. Causing invasive disease is an evolutionary dead end for these pathogens, so studies of carriage will not observe selection for variations that advantage bacteria in the blood or CSF. Current knowledge is mostly focused at the serotype and MLST level, and lacks the resolution and sample size to be able to address this question [26-28]. Though the only whole genome based study suggests this is not case (at the gene level) in *S. pneumoniae* [29], we believe higher powered study designs are needed to better answer this question.

We also hypothesise that bacterial variation may also occur during the invasive phase of meningitis. We have previously reported in a single patient that the bacteria appeared to adapt to the distinct conditions of blood and CSF [24]. These are very different niches from that of nasopharyngeal carriage where this variation is well documented [30], not least because the bacteria are under more intense exposure to immune pressures and have less time over which to accumulate mutations.

To look for adaptation to these three niches, we used samples from the MeninGene study [31, 32], based at the Academic Medical Centre Amsterdam. 938 patients recruited to the study had culture positive bacterial meningitis with samples collected from both their blood and CSF (breakdown by species in Table S1), and 54 with pneumococcal or meningococcal meningitis have a matched sample taken from their nasopharynx. By whole genome sequence analysis of large numbers of paired bacterial isolates cultured from these samples, we have been able to test for repeated variation that occurs during the course of the disease.

## Results

We made assumptions about the evolution of bacteria within the host, under which we discuss the power of pairwise comparisons between single colonies taken from each niche to capture repeated evolution occurring post-invasion:

1. There is a bottleneck of a single bacterium upon invasion into the first sterile niche (usually blood), which then founds the post-invasion population [7, 33].
2. A large invasive population is quickly established, as the population size approaches the carrying capacity of the blood/CSF. The population size is large enough for selection to operate efficiently.
3. As infection occurs in a mass transport system, populations are well mixed without any substructure. Therefore, the effective population size equals the census population size.
4. The bacterial growth curve within blood and CSF is similar.

Initially the population size is small, so selection is inefficient and the population-wide mutation rate is low. However, the eventual carrying capacity of the blood and CSF are large enough (>1.5×10^5^) [34, 35] for beneficial mutations to fix rapidly. Due to the short generation time of around an hour [36], this carrying capacity is reached early in the course of the disease (after 1-2 days) [37].

Crucially, population sizes where selection acts efficiently [38] are reached even earlier than this - a few hours after invasion. Therefore, mutations with a selective advantage occurring after the first stages of infection will eventually become fixed in the niche’s population. So, sequence comparison between colony picks from each niche is likely to find adaptation that has occurred post invasion.

Similarity of the bacterial growth curve within blood and CSF is an important assumption because in 45% of the pneumococcal cases there was evidence that CSF invasion happened before blood invasion (patients had a documented prior CSF leak, otitis media or sinusitis [39, 40]). This allows us to search for post-adaptation invasion that happens in either direction in this species. We investigated the validity of this assumption using analysis of data on the ivr locus (see Methods).

In carriage samples, although the population size is small [41] carriage episodes can persist over many months [42], therefore allowing the potential for mutations conferring an advantage in an invasive niche to arise. Additionally, during carriage there is known to be population wide diversity [30] and in some cases competition between strains [43]. We only sample a single strain from this diverse pool, which means we have less power to detect mutations either side of the bottleneck. Combined with the small sample size, this means only adaptive mutations with large selective advantages will be discovered in this part of the study. We therefore discuss our results from blood and CSF comparisons first, as we have a higher power to detect variation between these niches.

### No repeated post-invasion adaptation in coding regions across species

We performed comparisons between the pan-genome of each pair of blood and CSF isolates, using reference-free variant calling techniques (see Methods). The method was evaluated using simulated data, giving us confidence that it could detect the small amounts of variation expected between each isolate pair (figures S1, S2).

For each species we then counted the number of variants of any type between each blood/CSF isolate pair taken from a patient. For all species except *N. meningitidis* the majority of paired samples have no variation between them (Fig. 1).

**Fig 1:**
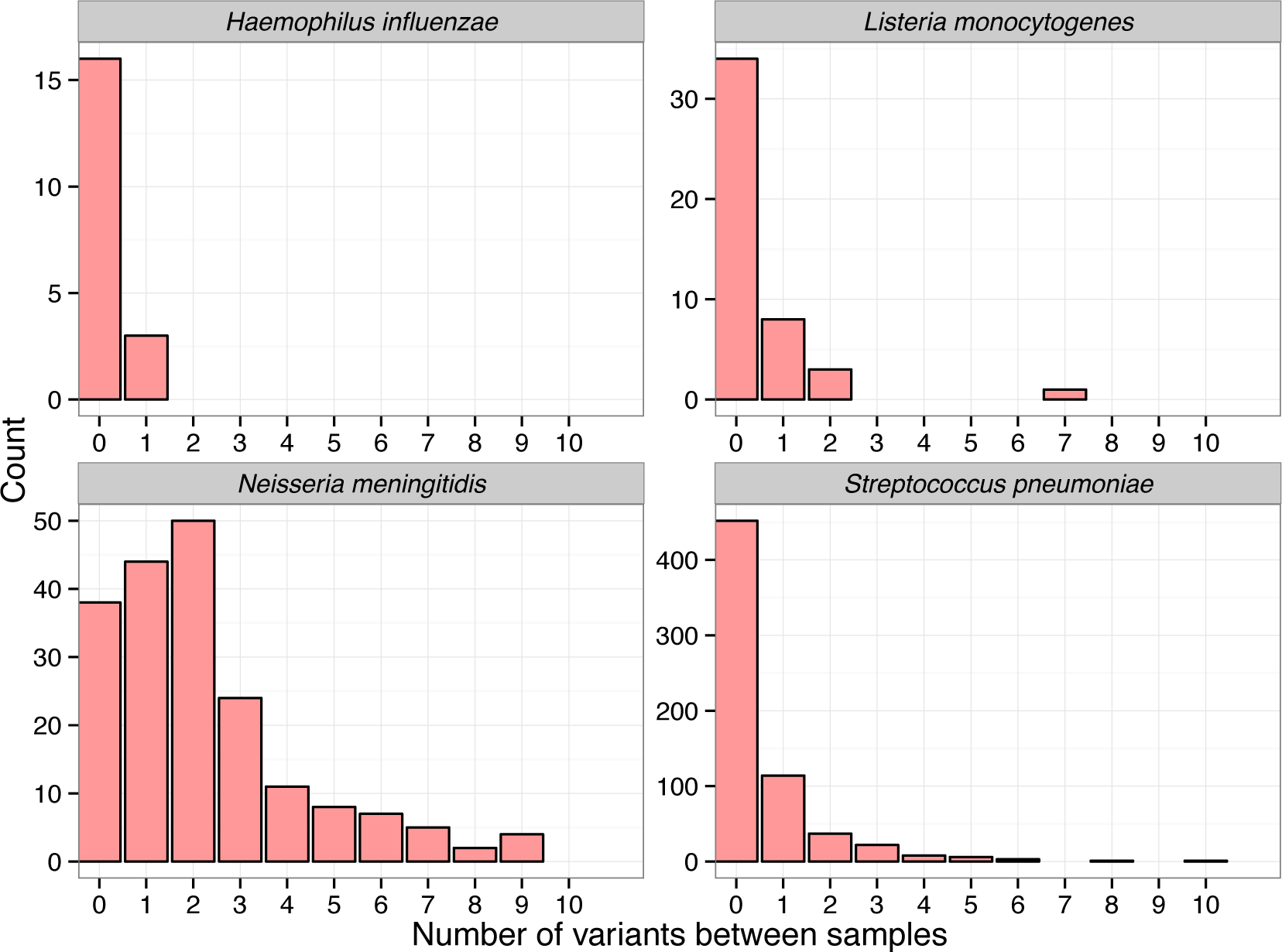
Histograms binned by number of variants between a blood/CSF sample pair, for each bacterial species. SNPs are from mapping, INDELs are from cortex. Three *S. pneumoniae* and one *N. meningitidis* sample with over 10 variants not shown.

In *S. pneumoniae* 452 of 674 paired samples (67%) were identical. The distribution is roughly Poisson (mean = 0.547), excluding outliers. In *H. influenzae* and *L. monocytogenes* the observed number of mutations between each paired set of strains is low, and similar in distribution to *S. pneumoniae*.

Variation between *N. meningitidis* pairs also followed a roughly Poisson distribution (mean = 2.34), which when compared to other species showed a higher number of variants between blood and CSF isolates (Wilcoxon rank-sum test, W = 25790, p-value < 10^−16^) such that most pairs have at least one variant between the blood and CSF samples.

In *N. meningitidis* these mutations may be a signal of repeated adaptation between the two niches if they cause the same functional change. Similarly, rare mutations in the other three species, if they cause the same functional change, could represent a signal of adaptation. To determine whether this is the case, we counted the number of times each gene annotation contained variation between the blood and CSF isolate over all the pairs collected, and used a Poisson test to determine whether this was more than expected for each gene (see Methods). For *L. monocytogenes* and *H. influenzae* there was not enough variation measured in the samples to show any such signal. For *S. pneumoniae* the results are shown in table 1.

**Table 1:** Genes containing significantly repeated mutations between blood and CSF isolate pairs in *S. pneumoniae*. Ordered by increasing p-value; locus tags refer to the D39 genome, if present.

| Gene name | Gene length (bp) | Mutations between blood and CSF | p-value |
| --- | --- | --- | --- |
| *pde1* (SPD_2032) | 1973 | 19 | $<10^{-10}$ |
| *dltD* (SPD_2002) | 1269 | 13 | $<10^{-10}$ |
| *dltB* (SPD_2004) | 1245 | 12 | $<10^{-10}$ |
| *dltA* (SPD_2005) | 1551 | 11 | $<10^{-10}$ |
| *clpX* (SPD_1399) | 1233 | 7 | $1.3 \times 10^{-8}$ |
| *wcaJ* (SPD_1620) | 693 | 6 | $3.4 \times 10^{-8}$ |
| *cysB* (SPD_0513) | 909 | 5 | $1.6 \times 10^{-5}$ |
| *cbpJ* | 1122 | 5 | $4.7 \times 10^{-5}$ |
| *amiC* (SPD_1670) | 1332 | 4 | $6.0 \times 10^{-3}$ |
| *marR* | 435 | 3 | $9.6 \times 10^{-3}$ |
| *fhuC* | 519 | 3 | $1.6 \times 10^{-2}$ |

The *dlt* operon, responsible for D-alanylation in teichoic acids in the cell wall [44-46], was the most frequently mutated region: 36 mutations in 31 sample pairs (Poisson test p<10^−10^).

Mapping the variation between sample pairs to the R6 *S. pneumoniae* strain, which has a functional *dlt* operon, the variations were annotated with their predicted function. There was no directionality to the mutations: 19 occurred in the blood, and 11 in the CSF. Only seven of the patients infected by these strains showed signs of blood invasion before CSF invasion (sinusitis or otitis); this also did not show directionality. The nature of the mutations is shown in Fig. 2 and Table S2. Most of these mutations would be expected to cause a loss of function (LoF) in the operon. Though this suggests this locus has a deleterious effect in invasive disease generally, the lack of directionality to the mutations means it does not show evidence of adaptation to either the blood or CSF specifically.

**Fig 2:**
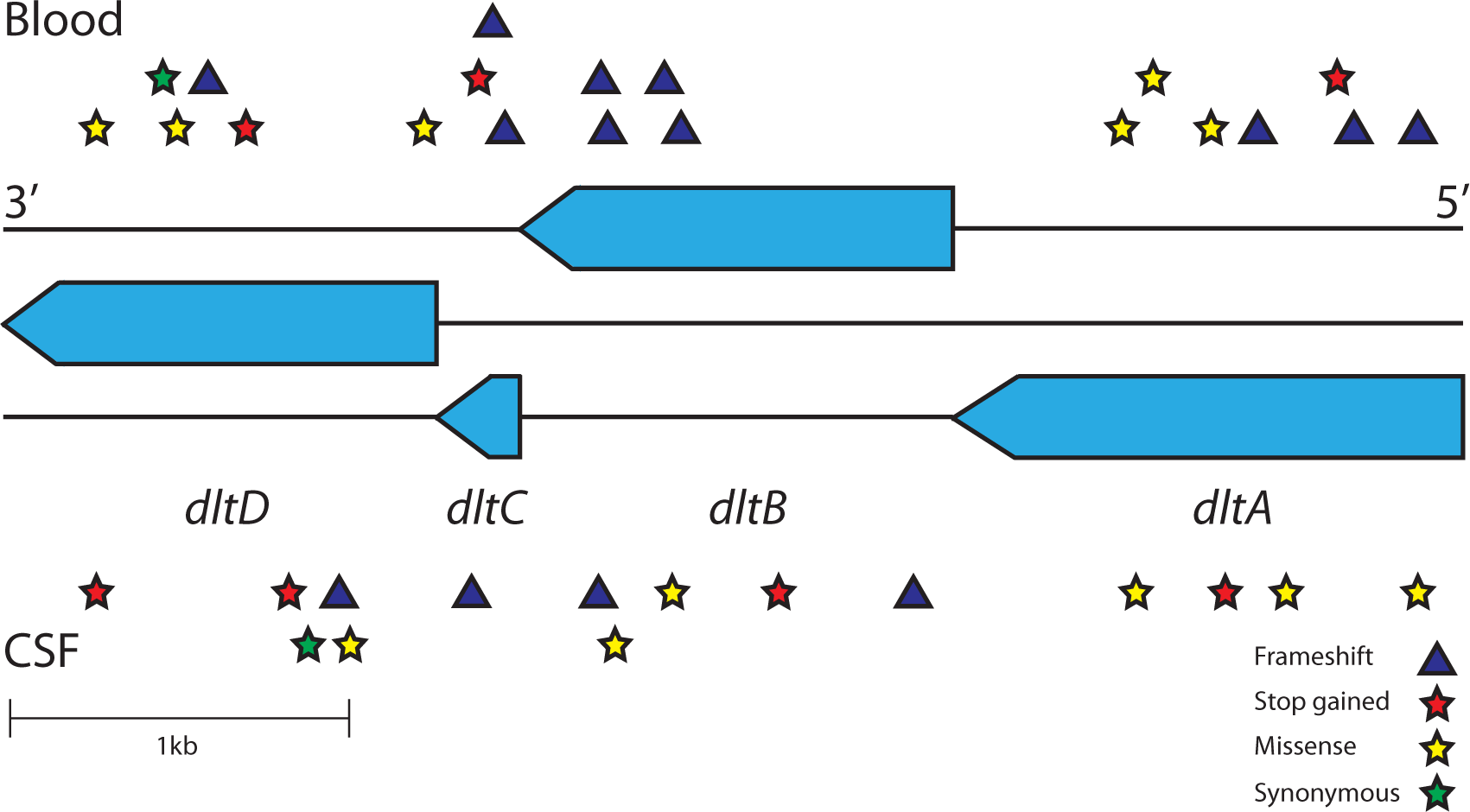
Mutations observed between all paired samples in the *dlt* operon. The operon consists of four genes in the three reading frames of the reverse strand. Mutations, displayed by type, in the blood strains are shown above the operon, and in the CSF strains below the operon.

In all the other genes in table 1 the variants are non-synonymous SNPs distributed evenly between blood and CSF, therefore also showing no adaptation specific to either niche.

The most frequently mutated genes between pairs in *N. meningitidis* are shown in table 2.

**Table 2:** Genes containing significantly repeated mutations between blood and CSF isolate pairs in *N. meningitidis*. Ordered by increasing p-value; locus tags refer to the MC58 genome, if present.

| Gene name | Gene length (bp) | Mutations between blood and CSF | p-value |
| --- | --- | --- | --- |
| *pilE* (NMB0018) | 384 | 18 | $<10^{-10}$ |
| *lgtC* | 189 | 16 | $<10^{-10}$ |
| *hyaD* | 327 | 14 | $<10^{-10}$ |
| *oatA* | 1869 | 19 | $<10^{-10}$ |
| *hpuB* (NMB1668) | 2382 | 17 | $<10^{-10}$ |
| *porA* (NMB1429) | 1178 | 12 | $<10^{-10}$ |
| *lgtA* (NMB1929) | 1050 | 10 | $<10^{-10}$ |
| *kfoC* | 360 | 7 | $<10^{-10}$ |
| *cotSA* | 1134 | 7 | $9.2 \times 10^{-9}$ |
| *ssaI* | 3252 | 6 | $3.9 \times 10^{-4}$ |

Top ranked are those relating to the pilus: *pilE* (19), *pilC* (6) and *pilQ* (4). pilE genes are associated with immune interaction [47], and are therefore expected to be under diversifying selection; an excess of non-synonymous mutations (46/49 observed; 33/49 expected for neutral selection) is consistent with this. The other notable gene with more mutations than expected in *N. meningitidis* was *porA*, a variable region which is a major determinant of immune reaction [48], in which 12 samples had frameshift mutations in one of two positions. Phase variation in the gene’s promoter region, affecting its expression, is discussed in more detail below.

The mutations in table 2 show no association with blood or CSF specifically, so do not represent adaptation to either niche. Genetic variation in *pilE, hpuA, wbpC, porA* and *lgtB* within host has been observed previously in a single patient with a hypermutating *N. meningitidis* infection [25]. The coding sequences with excess variants that are found in the samples analysed here include these genes. This also suggests an elevated background rate in these sequences, rather than strong selection between the blood and CSF niches.

### No evidence for repeated adaptation in intergenic regions in *S. pneumoniae* and *N. meningitidis*

Our previous result suggesting adaptation from blood to CSF was an intergenic change affecting the transcription the *patAB* genes, encoding an efflux pump. In general it is known that in pathogenic bacteria a common form of adaptation is mutation in intergenic regions, which may affect global transcription levels, causing a virulent phenotype [49, 50], antimicrobial resistance [51] and changing interaction with the host immune system [52]. Changes in these regions have previously been shown to display signs of adaptation during single cases of bacterial disease [18].

We therefore separately investigated the mutations in non-coding regions, which were only observed in *S. pneumoniae* and *N. meningitidis*. As genome annotations from the calling above are not consistent between samples outside of CDSs, we mapped the variation in intergenic regions to the coordinates of a reference genome. In a subset of samples we had carriage isolates corresponding to blood and CSF isolate pairs (discussed further below); we used these carriage isolates as the reference genome to determine whether these mutations occur in the blood or CSF isolate.

Figure S3a shows all mutations plotted genome-wide in *S. pneumoniae*. The peaks correspond to mutations in genes described in table 1. In the remaining 121 mutations in non-coding regions we observed no clustering by position. Over all pairs of samples, intergenic mutations were spread between blood and CSF isolates when compared to a carriage reference. This suggests none of the intergenic mutations are providing a selective advantage in either invasive niche.

The mutations in *N. meningitidis* are plotted in figure S4a, 110 of which were in non-coding regions. We observed enrichment, but no niche specificity, in the upstream region of six genes. These mutations are listed in table 3. Some of the mutations upstream of *porA* and *opc* are in phase variable homopolymeric tracts, which are discussed more fully in the section below. The other mutations are upstream of the adhesins *hsf*/NMB0992 and NMB1994, which are involved in colonisation [53] and immune interaction during invasion [54], and *frpB*/NMB1988 which is a surface antigen involved in iron uptake [55]. Differential expression of these genes may be an important factor affecting invasion, but the mutations we observed that may affect this do not appear to be specific to blood or CSF.

**Table 3:** Intergenic regions containing significantly repeated mutations between CSF and blood isolate pairs in *N. meningitidis*. Ordered by increasing number of mutations; coordinates refer to the MC58 genome.

| Coordinates | Downstream gene(s) | Mutations between blood and CSF |
| --- | --- | --- |
| 1468329-1468331 | *porA* (NMB1429) | 7 |
| 1072215-1072328 | *opc* (NMB1429) | 7 |
| 1008872-1008985 | *hsf* (NMB0992) | 6 |
| 1315621-1315672 | NMB1299 | 6 |
| 2092257-2092552 | *frpB* (NMB1988) | 5 |
| 2100124-2100258 | NMB1994 | 4 |

### No evidence for repeated adaptation in phase variable regions in *S. pneumoniae* and *N. meningitidis*

Phase variable regions, which may also be intergenic, can mutate rapidly and are known to be a significant source of variation in pathogenic bacteria [56]. This mutation is an important mechanism of adaptation [57], and meningococcal genomes in particular contain many of these elements [58].

In *N. meningitidis* we observed six samples with single base changes in length of the phase-variable homopolymeric tract in the *porA* gene’s promoter sequence, and five samples with the single base length changes in the analogous promoter sequence of *opc*. While changes in the length of these tracts will affect expression of the corresponding genes, both of which are major determinants of immune response [59, 60], the tract length does not correlate with blood or CSF specifically. *porA* expression has previously been found to be independent of whether isolates were taken from CSF, blood or throat [59].

In *S. pneumoniae*, recent publications highlight a potential role in virulence for the *ivr* locus, a type I restriction-modification system with a phase-variable specificity gene allele of *hsdS* in the host specificity domain (Fig. 3) [19, 20, 61]. There are six possible different alleles A-F (Figure S5) for *hsdS*, each corresponding to a different level of capsule expression. Some of these alleles are more successful in a murine model of invasion, whereas others are more successful in carriage. We used a mapping based approach (see methods) to determine whether any of these alleles were associated with either the blood or CSF niche specifically, which could be a sign of adaptation.

**Fig 3:**
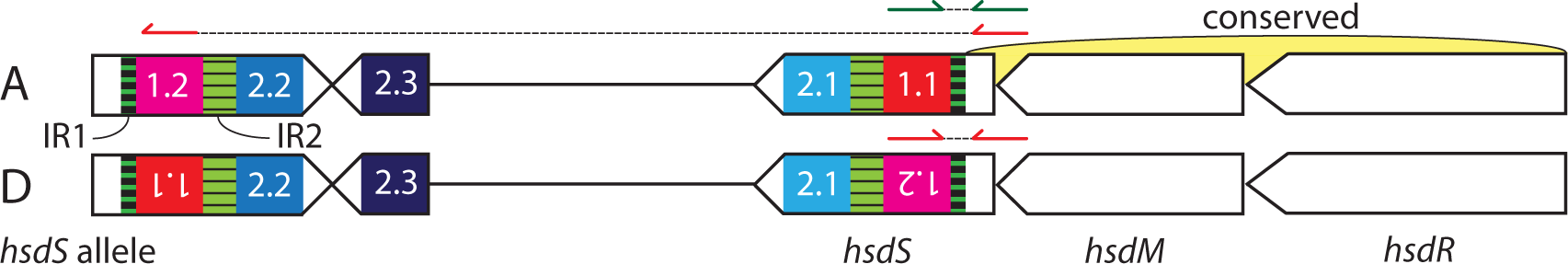
The structure of the *ivr* type I restriction-modification locus in *S. pneumoniae*. The restriction (*hsdR*) and methylation (*hsdM*) subunits, and the 5′ end of the specificity subunit (*hsdS*) are generally conserved. Inverted repeats IR1 (85bp) and IR2 (333bp) facilitate switching of downstream incomplete *hsdS* elements into the transcribed region. Top: The green read pair has the expected insert size, and suggests allele A (1.1, 2.1) is present. The red read pair is in the wrong orientation and has an anomalously large insert size. Bottom: The red read pair is consistent with the displayed inversion, suggesting allele D (1.2, 2.1) is present.

As the locus inversion is rapid and occurs within host, we first ensured that cultured samples are representative of the original clinical samples using PCR quantification of each allele. However, as even a single colony contains heterogeneity at this locus, simply taking the allele with the most reads mapping to it in each sample gives a poor estimate of the overall presence of each allele in the blood and CSF niches. To take into account the mix of alleles present in each sample, and to calculate confidence intervals, we developed a hierarchical Bayesian model for the *ivr* allele (see Fig S6 and Methods). This simultaneously estimates the proportion of each colony pick with alleles A-F for both each individual isolate (π), and summed over all the samples in each niche (μ). We apply this over *i* samples and *c* niches (in this case *c* can be blood or CSF).

For each pair of blood and CSF samples listed in Table S1, the difference in allele prevalence π_CSF_ − π_blood_ was calculated (table S3). All *S. pneumoniae* samples had a difference in mean of at least one allele (as the confidence intervals overlap zero), highlighting the speed at which this locus inverts.

While this means that between a single CSF and blood pair the allele at this locus usually changes, it is the mean of μ_*c*_ (corresponding to the mean allele frequency in each niche over all sample pairs) which tells us whether selection of an allele occurs in either the blood or CSF more generally. This is plotted in Fig. 4. As the confidence intervals overlap, no particular allele is associated with either blood or CSF *S. pneumoniae* isolates.

**Fig 4:**
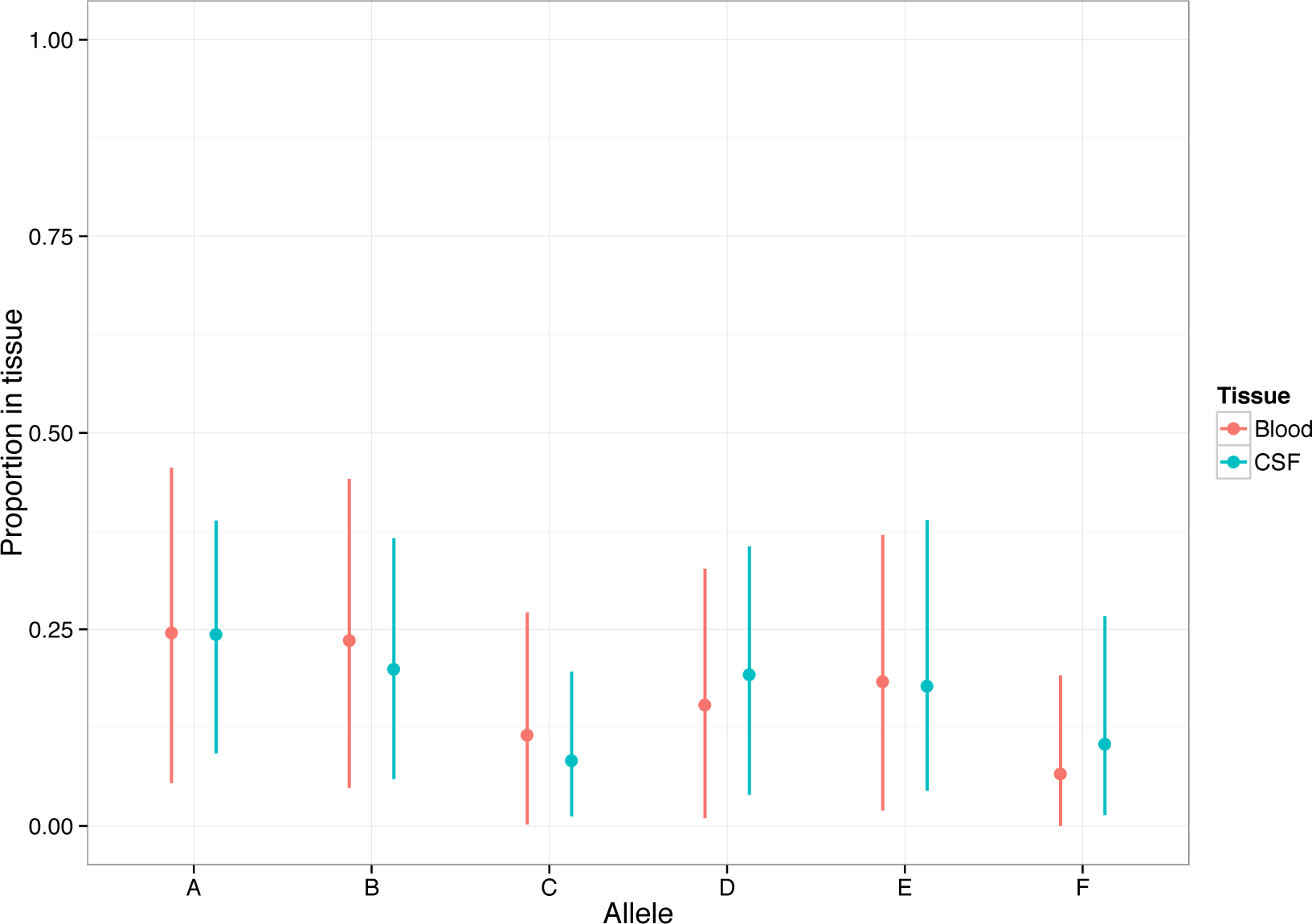
Mean and 95% highest posterior density (HPD) for *μ_c_*. This shows the proportion of each allele present in each of blood (red) and CSF (turquoise) tissues pooling across all samples.

Previous work on a murine invasion model [19] has shown an increase in proportion of alleles A and B over the course of infection. We did not observe the same effect in our clinical samples, though the large confidence intervals from the mathematical model suggest that genomic data with a small insert size relative to the size of repeats in the locus is limited in resolving changes in this allele. A small selective effect of *ivr* allele between these niches would therefore not be detected, but we can rule out strong selection for a particular allele (odds ratio > 2). Application to read data from a large carriage dataset may help resolve whether the same effect does occur in humans, as it would provide a greater temporal range over the course of pathogenesis.

### Carriage and invasive disease sample pairs show some evidence of repeated adaptation

Using the same methods, we also sequenced and analysed pairs of genomes from 54 patients that were collected from the nasopharynx and CSF. Six of these were *S. pneumoniae*. In these *S. pneumoniae* samples, we detected only one sample with any variation, which was a two base insertion upstream of the *gph* gene (Figure 5). This is similar to the amount of mutation observed between the blood and CSF isolates, which is expected given the similar sampling timeframes. While we found that a functional *dlt* operon appears to have a deleterious effect in invasive disease, we did not observe mutation between our carriage and disease samples. However, this was expected given the small number of carriage samples relative to the effect size detected for this operon.

**Fig 5:**
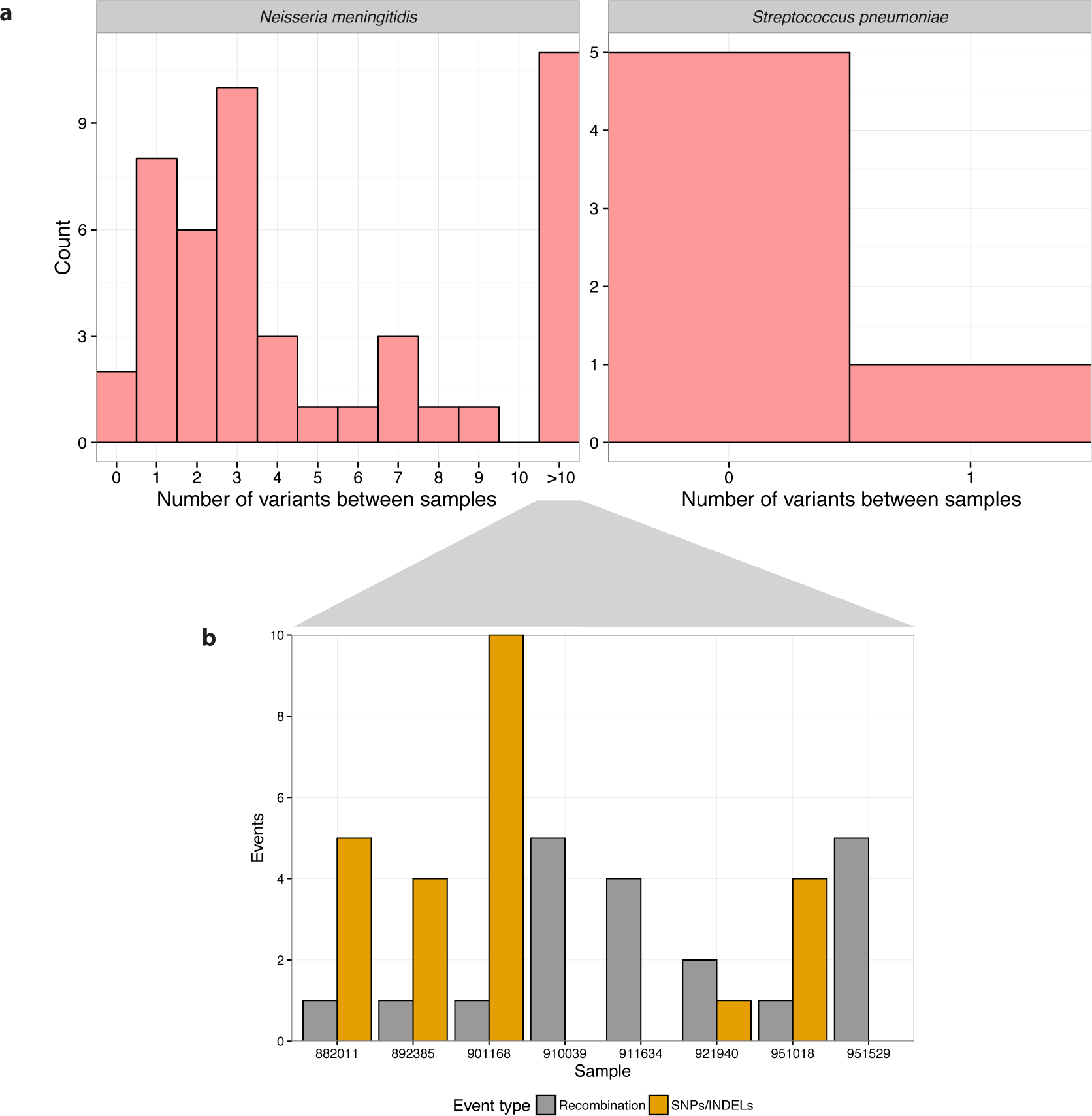
Histograms binned by number of variants between a carriage/CSF sample pair, for each bacterial species. a) As figure 1. In *N. meningitidis* eleven samples with over ten variants between them due to recombination events are grouped. b) The number of recombination and SNP/INDEL events in samples in the group with over ten detected variants

Between the 48 *N. meningitidis* carriage and CSF isolate pairs small numbers of mutations were common. We went on to search for regions enriched for mutation, however in 8 samples we observed large numbers of mutations clustered close together. These represent single recombination events, so when analysing genes enriched for mutation we counted each recombination as a single event [62, 63].

Table 4 shows the results of this analysis. Similar genes are mutated as in the blood/CSF pairs, again with no specificity to either niche. In phase variable intergenic regions, we observed four sample pairs with an insertion or deletion in the *porA* promoter tract with no niche specificity. Otherwise, none of the regions above showed enrichment for mutation in either niche. These observations support the theory that these genes mutate at a higher rate but do not confer a selective advantage in any of the three niches studied.

**Table 4:** Genes containing significantly repeated mutations between nasopharyngeal and CSF isolate pairs in *N. meningitidis*. Ordered by increasing p-value; locus tags refer to the MC58 genome, if present.

| Gene name | Gene length (bp) | Mutations between nasopharynx and CSF | p-value |
| --- | --- | --- | --- |
| *lgtA* (NMB1929) | 1050 | 6 | $5.0 \times 10^{-7}$ |
| *oatA* | 1869 | 6 | $1.5 \times 10^{-5}$ |
| *hyaD* | 327 | 4 | $2.6 \times 10^{-5}$ |
| *pilE* (NMB0018) | 384 | 4 | $3.8 \times 10^{-3}$ |
| *pilT* (NMB0052) | 1131 | 4 | $3.5 \times 10^{-3}$ |
| *dca* (NMB0415) | 444 | 3 | $1.1 \times 10^{-2}$ |

A notable exception to this is the *dca* gene, a phase variable gene involved in competence in *Neisseria gonorrhoea* but of unknown function in *N. meningitidis* [64, 65], in which all mutations are protein truncating variants in the invasive isolate. Similarly, though not reaching significance (due to the long length of the genes) were the *ggt* (NMB1057) and *czcD* (NMB1732) genes in which three recombinations occurred, all of which were in the invasive isolate of the pair.

The mutations in these three genes therefore may confer a selective advantage in the invasive niche; the sequence at these loci in the invasive strains are the same as the MC58 reference, an invasive isolate itself.*ggt* has previously shown to be essential for *N. meningitidis* growth in CSF in rats [66], and metal exporters such as *czcD* have been shown to increase virulence in a mouse sepsis model [67]. More such paired carriage and invasion samples would be needed to confirm this is the case in human invasive disease.

## Discussion

Previous studies have shown that substantial levels of genomic DNA sequence variation occur in bacteria colonising or infecting human hosts [12, 14, 15] and suggest that some of this variation may be due to selective adaptation [18, 22-24, 68]. Such adaptations during invasive bacterial disease could lead to new insights into the processes of pathogenesis with the potential to inform therapies[69, 70]. Therefore it is important to capture evidence of such variation; we sought to do this through large-scale genomic analysis. We have searched for variation in four different bacterial species comparing genomes from bacteria isolated from both blood and CSF from the same individuals in 938 bacterial meningitis cases.

We found overall that the amount of variation within bacteria that occurs after infection is low. The mutation observed is not randomly distributed throughout the genome, but is randomly distributed between blood and CSF isolates. These mutations are therefore an observation of a higher mutation rate in these regions during invasion (for example the pilus in *N. meningitidis*, which is known to be under diversifying selection) but not repeated adaptation to either niche. This study indicates that our previous observation of variation between blood and CSF isolates from a single case of meningitis was a rare event most likely driven by antibiotic selection pressure during treatment.

We went on to analyse 54 samples comparing carriage and invasive isolates from the same patient. Though the sample size was lower, and we did not fully sample diversity within the nasopharynx, we were able to get an insight into potential genetic differences between bacteria in these niches. We see some of the same genes mutate rapidly between blood and CSF isolates also doing this between carriage and invasion. This supports the conclusion that these genes have a higher mutation rate, rather than giving a selective advantage to a niche. Finally, we saw possible evidence for selection on the *dca* gene in *N. meningitidis*.

This study eliminates the need to search for bacterial diversity between invaded host niches (blood and CSF) when trying to explain pathogenesis of meningitis. However, our comparison between the genomes of carriage and invasive isolates did show some weak signals of adaptation. Our power in these comparisons was limited by small sample size and single colony sequencing. Future studies are needed to comprehensively identify whether adaptation occurs through bacterial genetics between carriage and invasion. This should be addressed through either comparing more pairs of carriage and invasive isolates with deeper sequencing, or large scale genome wide association studies.

## Methods

### Ethics statement

Written informed consent was obtained from all patients or their legally authorized representatives. The studies were approved by the Medical Ethics Committee of the Academic Medical Center, Amsterdam, The Netherlands (approval number: NL43784.018.13).

### Reference free variant calling

We extracted DNA from positive blood and CSF cultures from adults with bacterial meningitis in the Netherlands from 2006 to 2012, and sequenced this with 100bp paired end reads on the Illumina HiSeq platform.

To avoid reference bias, and missing variants in regions not present in an arbitrarily chosen reference genome, we then performed reference free variant calling between all sequence pairs of isolates using two methods: the ‘hybrid’ method [71] and Cortex [72]. The former uses *de novo* assembly of the CSF sequence reads, mapping of reads from both the blood and CSF samples back to this sequence, then calling variants based on this mapping. Cortex uses an assembly method that keeps track of variation between samples as it traverses the de Bruijn graph.

In the hybrid method we created draft assemblies of the CSF samples using SPAdes v3.5 [73], corrected these with SSpace and GapFiller [74], then mapped reads from both blood and CSF samples to this reference using SNAP [75] followed by variant calling with bcftools v1.1 [76].

For Cortex we first error corrected sample reads using quake [77], preventing false positive calls supported by very low coverage of reads. The joint workflow of cortex was then used with each set of corrected reads in its own path in the de Bruijn graph, and bubble calling was used to produce a second set of variants between samples. SNPs in the error corrected reads were also called using the graph-diff mode of SGA [78].

### Simulations of closely related genomes

As the rate of variation is very low, we needed to ensure we had sufficient power to call variants and didn’t suffer from an elevated false negative rate. We did this by simulating evolution of *S. pneumoniae* genomes along the branch of the tree between *S. pneumoniae* R6 [79] and the common ancestor with *Streptococcus* mitis B6 [80] (figure S7). The rates in the GTR matrix and insertion/deletion frequency distributions were estimated by aligning the R6 and B6 reference sequences with Progressive Cactus [81]. An average of 200 mutations with these rates were created in 100 sequences, and Illumina paired end read data at 200x coverage simulated using pIRS [82]. Variants between these sequences and a draft R6 assembly from simulated read data were then called using both of the above methods; comparison with the mutations known to be introduced allowed power and false positive rate to be calculated - separately for SNPs (single base substitution) and INDELs (one or more bases inserted or deleted).

In addition to *in silico* simulation, we cultured blood/CSF paired strains 4038 and 4039 [24] and resequenced them using the same 100bp Illumina paired end sequencing as the rest of the isolates in the study. The genomes of strains 4038 and 4039 have been exhaustively analysed using multiple sequencing technologies, so represent high quality positive control data to assess the calling methods. Both methods were tested on these data.

The highest power was achieved using hybrid mapping for SNPs and Cortex for INDELs: median power for calling SNPs was 90% using hybrid mapping, and 74% for INDELs using cortex (figures S1 and S2). This combination of methods, mapping for SNPs and cortex for INDELs, was therefore used across all samples. When applied to the paired strains 4038/4039 the same mutations as originally reported are recovered, plus a 37bp insertion in *cysB* which was found to be introduced during culturing.

We used simulations to compare against a simple method of mapping against an arbitrary reference, in this case TIGR4 [83]. We found our reference free method has greater power, especially for INDELs (figure S8), and a markedly reduced false positive rate. We also used an assembly method alone to compare gene presence and absence, but this too suffered from a vastly elevated false positive rate (figure S9).

### Tests for genes, intergenic regions and genotypes enriched for mutation

To scan for repeated variation, the number of mutations in each CDS annotation (adjusted for CDS length) was counted. We then performed a single-tailed Poisson test using the genome wide mutation rate per base pair multiplied by the gene length as the expected number of mutations. The resulting p-values we corrected for multiple testing using a Bonferroni correction with the total number of genes tested as the *m* tests. Tables 1 and 2 report those CDS with an adjusted p-value less than the significance level of 0.05. For intergenic regions, anywhere with more than one variant is reported.

To test whether certain genotypic backgrounds are associated with a higher number of mutations that occurs post-invasion, a linear fit of each MLST against number of mutations between blood and CSF isolates was performed. The p-values of the slope for each MLST were Bonferroni corrected; at a significance level of 0.05 no MLST was associated with an increased number of mutations. For genes the same test was performed, except samples were coded as one and zero based on whether they had a mutation in the gene being tested or not. We performed a logistic regression for each gene with over ten mutations in tables 1 and 2: no genes being mutated post invasion were associated with an MLST.

### Copy number variation

We called copy number variations (CNVs) between samples by first mapping each species to a single reference genome, then fitting the coverage of mapped reads with a mixture of Poisson distributions [84]. Regions were ranked by the number of sample pairs containing a discordant CNV call. As with regions of enhanced SNP/INDEL variation, we performed a Poisson test for enrichment on these regions.

In *S. pneumoniae* the most frequently varying region was due to poor quality mapping of a prophage region. The only other region with p < 0.05 was a change in copy number of 23S rRNA seen in a small number of sample pairs. In *N. meningitidis* mismapping in the *pilE/pilS* region accounts for the only CNV change. No CNV changes were observed in multiple sample pairs of either *L. monocytogenes* or *H. influenzae*.

### Variant annotation

To then be able to compare between samples using a consistent annotation, we mapped the called variants to the *S. pneumoniae* ATCC 700669 reference [85] for *S. pneumoniae*, and MC58 [86] for *N. meningitidis*. This was done by taking a 300 base window around each variant and using blastn on these with the reference sequence. We used VEP [87] to annotate consequences of each variant. The variants were visualised by plotting variant start positions between all pairs against a reference genome, which allowed identification of clustering of variation in unannotated regions.

### *ivr* locus allele determination directly measured from clinical samples

As the rate of variation at this locus is fast compared to other types of variation measured, we checked that culturing the bacteria does not cause the allele to change from what is observed in the original clinical sample. We therefore extracted DNA from a subset of 53 of 674 paired clinical CSF samples and the respective bacterial isolates.

Allele prevalence was quantified using a combined nested PCR protocol based on PCR amplification of the ivr locus, digesting with DraI and PleI followed by quantitative analysis of banding patterns on a capillary electrophoresis [19]. Allele prevalence was identical between the original clinical sample and cultured bacteria in 50 out of the 53 samples. The predictive power of the *in vitro* detected *ivr* allele prevalence in a pneumococcal culture for the original allele prevalence within the clinical sample is therefore sufficient to draw conclusions from.

### *ivr* locus allele determination from genomic data

There are six possible alleles A-F at this locus, though due to the high variation rate and structural rearrangement mediating the change the allele cannot reliably be determined using assembly and/or standard mapping of short read data.

Instead, mates of reads mapping to the reverse strand of the conserved 5' region were extracted for each sample, and mapped with BLAT [88] to the possible alleles in position 1. This forms a vector *r_i_* of length two for each sample *i*, with the number of reads mapped to 1.1 and 1.2. Similarly, to determine the 3′ allele (position 2), pairs of reads mapping to each of the reverse strand of allele 1.1 and the forward strand of allele 1.2 were extracted and mapped to the three possible alleles in position 2 (figure S5). This forms a vector *q*_*i*_ of length six for each sample *i*, with the number of reads mapped to each allele A-F.

From mapping, we found 621 sample pairs had at least one read mapping to an allele of the *ivr* locus *hsdS* gene (table S4). Those without any reads mapping had either a deletion of one component of the locus, or a large insertion mediated by the *ivr* recombinase.

### Bayesian model for *ivr* allele

We first modelled the state of the 5′ allele (TRD1.*j*) only. For the two possible alleles 1.1 and 1.2, the number of reads mapping to each allele (a 2-vector *r*_*i*_) was used as the number of successes in multinomial distribution *z*_*c*_ (*c* - index for niche). From these we infered the proportion of each allele in each individual sample π_*i*_, and in each niche overall μ_c_. This was done by defining Dirichlet priors expressing the expected proportion of an allele in a given sample π_i_ to be drawn from a Dirichlet hyperprior representing the proportion of the allele that is found in each niche as a whole μ_c_. The κ parameter sets the variance of all the individual sample allele distributions μ_ic_ about the tissue average μ_c_, with a higher κ corresponding to a smaller variance. This model is represented in figure S6.

The hyperparameter *A*_μ_ which encodes the total proportion of each allele we expected to see over all samples, was set to the average amount of the allele observed from the long range PCR in a subset of 53 paired samples, as described above.

The observed number of reads mapping to each allele, prior distributions defined above, and structure of the model in figure S6 defines a likelihood which can be used to infer the most likely values of the parameters of interest π and μ. We used Rjags to perform MCMC sampling to simulate the posterior distribution of these parameters. We used 3 different starting points, and took and discarded 30000 burn in steps, followed by 45000 sampling steps. Noticeable auto-correlation was seen between consecutive samples, so only every third step in the chain was kept in sampling from the posterior. We manually inspected plots of each hyperparameter value and mean at each point in the chain, as well as the Gelman and Rubin convergence diagnostic, which showed that the chain had converged over the sampling interval (figure S10).

To model both the 5′ end (TRD 1.1 and 1.2) and the 3′ end (TRD 2.1, 2.2 and 2.3) together, so each isolate *i* is represented by an allele A-F, for each isolate the total number of reads mapping *n*_*i*_ was drawn from the distribution in equation (1)

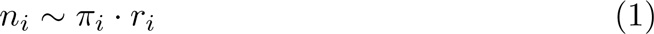

where *j* is the index of the TRD region, *r*_*ij*_ is the number of reads in sample *i* that had a mate pair downstream from TRD1.j mapping to any TRD2 region, and μ_i_ is the posterior for allele frequency in the sample.

The distribution for the number of reads mapping to each allele *j*, *z*_*ij*_ was defined as in equation (2)

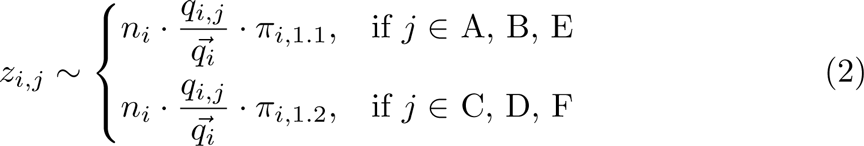

where *q*_*i*_ is a vector of length six which contains the number of reads mapped to each allele A-F as described above, and π, *i* and *n* are as previously. A single sample for *z* was taken for each isolate *i*. This 6-vector *z*_*ij*_ is then used as the observed data in the same model as above to infer π_i_, and μ_c_ for the whole locus allele (A-F) rather than just the 5′ end.

For the 5′ allele (TRD1.*j*) a model using a single κ parameter rather than a κ indexed by tissue *c* was preferred (change in deviance information criterion [89] ΔDIC = −0.523). For the 3′ allele (TRD2.*j*), a model with a single κ parameter did not converge. A model with κ indexed by allele was used instead.

### Diversity of *ivr* allele within samples

As the speed of inversion is rapid, the subsequent polymorphism of this locus was also used to evaluate our assumptions about diversity of the bacterial population within each niche. We calculated the Shannon index of each sample’s vectors π_CSF_ and π_blood_ to measure diversity of the sample in each niche. The mean Shannon index across CSF samples was 1.01 (95% highest posterior density (HPD) 0.39-1.51) and 0.98 (95% HPD 0.35-1.55) in the blood. Looking at each sample pair individually, the difference between diversity in each niche appears normally distributed with a mean of zero (figure S11). Together, these observations suggest a similar rate of diversity generation in each niche. This is in line with our assumption that the two populations have similar mutation rates, and a similar number of generations between being founded and being sampled.

## Acknowledgements

Thanks to Win Kit Man for DNA isolation of bacteria. Simon Harris and James Hadfield developed the visualization tool used plot variants across the genome (http://jameshadfield.github.io/phandango/). The study benefitted from collaboration in ESGIB.

### Funding

Work at the Wellcome Trust Sanger Institute was supported by Wellcome Trust core funding (098051 https://wellcome.ac.uk/). J.A.L. is supported by a Medical Research Council studentship grant (1365620 http://www.mrc.ac.uk/). This work was also supported by grants from the European Research Council (ERC Starting Grant [proposal/contract 281156] https://erc.europa.eu/) and Netherlands Organization for Health Research and Development (ZonMw; NWO-Vidi grant 2010 [proposal/contract 016.116.358] http://www.zonmw.nl/), both to D.v.d.B. The Netherlands Reference Laboratory for Bacterial Meningitis is supported by the National Institute for Health and Environmental Protection, Bilthoven (http://www.rivm.nl). NJC is funded by a Sir Henry Dale Fellowship, jointly funded by the Wellcome Trust and Royal Society (Grant Number 104169/Z/14/Z https://wellcome.ac.uk/, https://royalsociety.org/). MRO was supported by Medical Research Council grant (MR/M003078/1 http://www.mrc.ac.uk/).

